# Modelling *G×E* with historical weather information improves genomic prediction in new environments

**DOI:** 10.1101/213231

**Authors:** Jussi Gillberg, Pekka Marttinen, Hiroshi Mamitsuka, Samuel Kaski

**Affiliations:** Helsinki Institute for Information Technology HIIT, Department of Computer Science, Aalto University, PO Box 15400, 00076 Aalto, Finland.; Institute for Chemical Research, Kyoto University, Gokasho, Uji 6110011, Japan

## Abstract

Interaction between the genotype and the environment (*G×E*) has a strong impact on the yield of major crop plants. Although influential, taking *G×E* explictily into account in plant breeding has remained difficult. Recently *G×E* has been predicted from environmental and genomic covariates, but existing works have not shown that generalization to new environments and years without access to in-season data is possible and practical applicability remains unclear. Using data from a Barley breeding program in Finland, we construct an in-silico experiment to study the viability of *G×E* prediction under practical constraints. We show that the response to the environment of a new generation of untested Barley cultivars can be predicted in new locations and years using genomic data, machine learning and historical weather observations for the new locations. Our results highlight the need for models of *G×E*: non-linear effects clearly dominate linear ones and the interaction between the soil type and daily rain is identified as the main driver for *G×E* for Barley in Finland. Our study implies that genomic selection can be used to capture the yield potential in *G×E* effects for future growth seasons, providing a possible means to achieve yield improvements, needed for feeding the growing population.

---

Global yield improvements are needed to feed the growing population^1^. One possibility is to breed varieties for higher environmental adaptability, known as targeted breeding^2^. By improving the genetic fit of varieties in their growth environments, yield potential in the interaction between the genotype and environment could be realised. While the importance of *G×E* for agronomic performance is widely accepted, utilisation calls for methods that predict yields in new environments, because actual experimental data, consisting of yields of plant variety candidates from yield trials, will in practice be available from only a very limited number of environments. Importantly, prediction of a plant’s response to a new environment cannot be based on weather data from the growth season, as those will never be available at the time of prediction.

Methods for “cold start” prediction problems^3^, where predictions are needed for completely novel instances, have been developed within the machine learning community. Example applications include design of novel drugs for previously unseen cancers^4^, and recommendations in on-line shopping for new customers and/or products^3^. These methods are based on using external covariate data that describe properties of the novel instances. We develop a new method, an extension of the Kernelized Bayesian Matrix Factorization^5^, to account for the uncertainty in the covariates, which allows the use of historical records to predict weather conditions for future growth seasons, and eventually makes future *G×E* prediction for yield possible. Therefore, our new method, unlike the existing alternatives^6–8^, does not rely on accurate weather information from the growth season from the new location (Figure 1).

**Figure 1:**
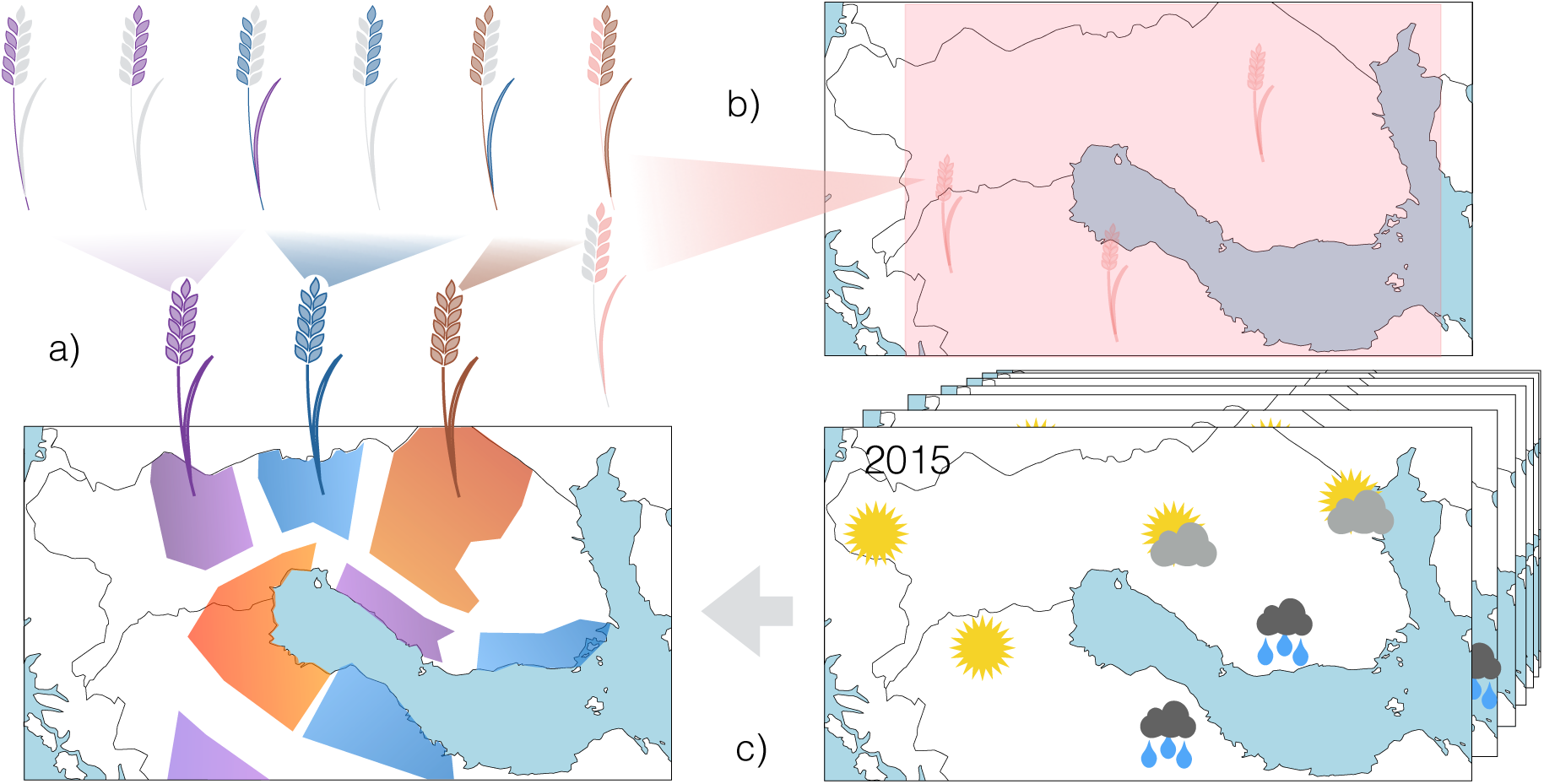
Outline of our approach. a) Precision breeding aims at producing varieties that are optimal for a specific environment. As compared to traditional breeding (b), targeted breeding aims at higher environmental adaptation, i.e., smaller target environments. Weather (microclimate) is a crucial driver for agronomic performance, but as it is unknown for future growth seasons, we use historical weather records (c) to predict the environmental stresses. The growth locations differ with respect to their estimated probabilities of extreme conditions and our method can be used to manage risks by trading-off yield potential for stress tolerance, when the risk in a particular environment is elevated.

In genomic selection (GS)^9^, field trials are replaced with genomic predictions to speed-up plant breeding. We formulate an *in silico* experimental setup for GS in targeted breeding that, unlike existing works^7, 8, 10–12^, strictly satisfies all realistic constraints: test locations, years, and genotypes are all genuinely new (not part of the training set) and yields are predicted for the offspring of the training set. In this setup, we demonstrate the feasibility of targeted breeding by investigating the accuracy of *G×E* prediction using environmental data including historical weather information but without in-season data 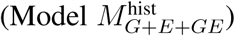. We compare this with multiple competing settings, including the non-realistic ideal situation having in-season data (*M*_*G+E+GE*_), a model without the *G×E* interaction (*M*_*G+E*_), a previous implementation with *G×E* interactions using in-season data (`GE-BLUP`)^8^, and the industry-standard that does not include *G×E* (best linear unbiased prediction using genomic data^13^; `GBLUP`). Data from a barley breeding program in Finland from Boreal Plant Breeding Ltd, including historical weather information for the target environments, are divided into training, validation, and test sets, and the prediction accuracy is measured as the average correlation between predicted and observed yields in the test sets^8^(Figure 2c). A sensitivity analysis is done to explore the impact of model assumptions. A description of the model and the setup can be found in Materials and Methods, and further details are given in the Supporting Information.

**Figure 2:**
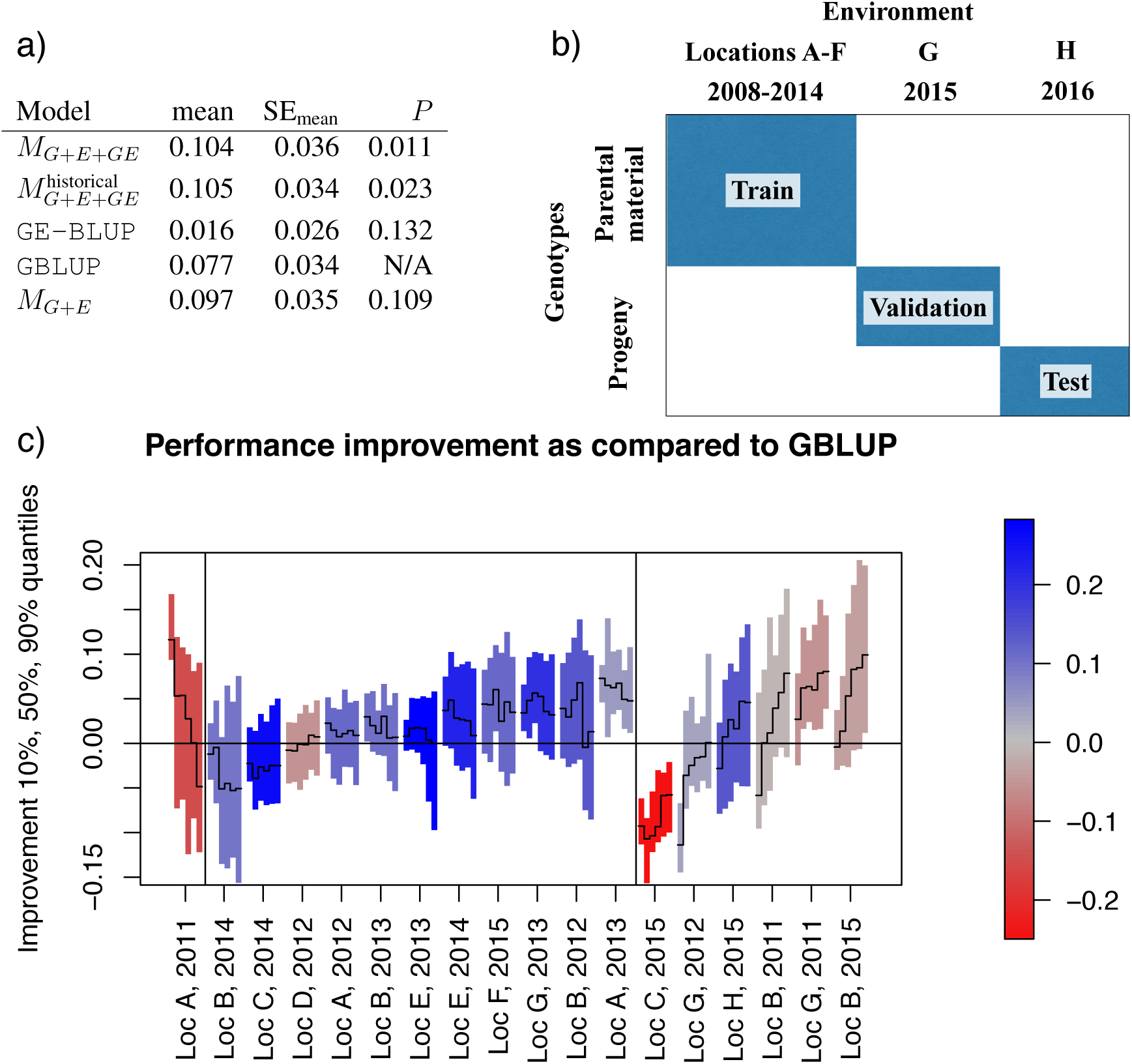
Predicting *G* x*E* with historical weather information improves genomic prediction accuracy in strictly new environments. a) Comparison of prediction accuracies; mean: correlation between predicted and observed yields, averaged across test environments; SE_mean_: standard error of the mean; P: p-value compared to the industry standard (`GBLUP`). *b*) Outline of the in silico setup for comparing methods. c) Sensitivity analysis: the difference in prediction accuracies (y-axis) between *G* xE prediction with historical data 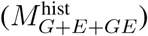 and the industry standard (`GBLUP`) is shown in 18 different environments (x-axis); values above the horizontal line mean that 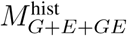 is mote accurate. Six vertical bars are shown for each environment, representing variability in results (median and 90 % confidence intervals). Starting from the left, they correspond to models with 0, 1, 2, 3, 4 or 5 *G×E* interaction terms (0 corresponds to the *M*_*G+E*_ model). The color indicates the performance of `GBLUP` in the environment, red meaning `GBLUP` performed poorly (Loc C, 2015 were omitted from the comparison as all methods performed poorly there). Vertical lines divide the environments into three groups: *G×E decreased*: including *G×E* terms to the model decreased performance; *G×E neutral*: 10 environments where *G×E* terms had neutral effect; *G×E* increased: 6 environments where performance increased by adding more *G* x*E* terms.

Modelling *G×E* with historical weather data, 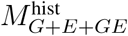, improves predictive accuracy as compared to the industry-standard, `GBLUP` (Figure 2a; p=0.011, a two-sided paired Wilcoxon signed rank test, df=17). The improvement is comparable to using in-season data (*M*_*G+E+GE*_, p=0.023). The Bayesian methods in general show higher accuracy whereas GE-BLUP performs poorly with the data available. Overall, the absolute prediction accuracy of all methods was relatively low in this challenging setup, with 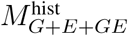 having the highest correlation of 0.105. Nevertheless, the improvement is considerable over the industry-standard with correlation 0.077, with the proposed new method explaining 85% more of the variation of the phenotype on average.

The sensitivity analysis demonstrates considerable variability between test environments (Figure 2c). Indeed, including *G×E* interaction terms into the model decreased accuracy in 1/18 environments, had little effect in 11/18 environments, but improved the accuracy substantially in 6/18 environments. In the last group, increasing model complexity by adding more *G×E* components consistently improved performance, which highlights the potential to increase accuracy through complex modelling of *G×E*. Importance of different data sources to the predictions can be further analysed by investigating the weights of the different kernels, used to summarise the data sources (Figure S 3). We see that the two most influential kernels were the ones that represented 1) the non-linear interaction between soil type and daily rainfall, and 2) the non-linear effect of rain, matching well the biological understanding of the problem.

Our experiments confirmed that prediction in new environments is a challenging task, as reported earlier^8^, our method reaching the highest correlation of 0.105 between predictions and observations. Nevertheless, the usefulness of including multiple *G×E* interaction terms and nonlinear interactions between environmental covariates became evident from our results. We expect that gains from modelling *G×E* will increase in the future as more data, representing further locations and years, will allow more accurately distinguishing the interactions from the main effects. Other ways to improve the predictions inlcude using more detailed genomic modeling, e.g. using Gaussian and other kernels for summarizing the SNP data.

Besides targeted breeding, there are several other needs for *G×E* prediction models. They could mitigate the problems of conventional breeding: accounting for historical weather in the actual target population of environments can help prevent overfitting to the conditions in the few field trials performed, as discussed in detail in SI Gains from modelling *G×E* for current target population of environments. The assumption of the match between field trials and actual growing locations is equally crucial for the official variety trials for value of cultivation and use (VCU), required in most countries to evaluate new varieties. *G×E* models are also needed in assessing the effects of climate change and to select for varieties that react favourably to the altering conditions^1^. For this purpose, the historical weather observations in 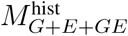 can be replaced with climate simulations to assess the performance of varieties under various climate scenarios. To summarise, we showed that *G×E* prediction in the setup required by targeted breeding, where the environments are strictly new and predictions are based on historical weather data available at the time of prediction, improves prediction accuracy significantly compared to the industry standard, which is needed to accelerate the implementation of targeted breeding.

## Methods

### Data

All data used in the experiment come from a barley breeding program in Finland, which is a part of a larger population of target environments for barley as varieties used in Finland are also used in other Nordic countries. The phenotype consists of (z-transformed) yield measurements (kg/ha) for 2,244 lines observed in trials at 11 locations across the 4 southernmost growth zones in Finland from 2008 to 2015. The total number of observed location x year combinations is 35. In some locations, trials have been performed on several years and several fields with varying soil properties, and a total of 12,277 yield observations have been recorded. The number of observations per genetic line ranges from 1 to 118 (median 4). The lines were genotyped with the Illumina 9k iSelect SNP Chip, SNPs with minor allele frequence (MAF) < 0.05 or with > 5% values missing were omitted. Also all genotypes with > 5% of SNPs missing were omitted. The final proportion of missing genotype data is 0.002.

The soil characteristics for each field block are measured in terms of the proportions of sand, silt and clay (soil classification triangle^14^) and the proportion of organic content. Meteorological information consists of daily averages of temperature and rainfall, and the distances to the closest meteorological station range from 1 to 40 km (average 13.5 km). The baseline approach GE-BLUP^8^ requires summarising the weather information per crop stage: vegetative (from sowing to visible awns), heading time (from visible awns to the end of anthesis), and grain filling (from the end of anthesis to maturity). The times of the crop stages are estimated using temperature sum accumulation; the details are given in Section Comparison methods. In the weather observations, the proportion of missing values in daily average temperature and rainfall measurements is < 0.0015 (max 3 missing values/environment) and < 0.0032 (max 2 missing values/environment), respectively.

### Experimental setup

To study prediction accuracy, we use a setup that strictly imposes the realistic constraints related to modelling *G×E* in targeted breeding for new locations. Predictions are required for new locations (not part of the experimental grid) and for years for which no phenotype data are available (to mimic future growth seasons). Additionally, predictions are needed for the offspring of the lines in the training set, which have no phenotype data observations. More details with a summary of the differences between our setup and earlier works are given in SI Details of experimental setup. We measure prediction accuracy using cross-validation, where the training, validation and test sets are selected to enforce the realistic constraints (Figure 2c). As the performance measure for prediction accuracy, we follow the conventional approach, i.e., the Pearson correlation between the predicted and observed yields in the test set^8, 10–12^. This correlation is computed for each cross-validation fold in turn, and averaged over the test cases. Similarly to Malosetti et al.^8^, the test case-specific correlations are transformed into Fisher’s z-scores before averaging and back-transformed to obtain the final results. We regress the *G×E* interactions on the average characteristics of the growing season: for each yield trial, we use the weather observations from the typical growing season (1^st^ of May until end of August) regardless of the sowing date. This indirect approach allows predicting with historical weather data. When predicting with historical data, the prediction for each genotype is made for each year for which historical weather observations are available, and the median of those is used as the final predicted value.

We also carry out a sensitivity analysis that allows studying the impact of modelling assumptions, such as inclusion of *G×E* interaction components to the model. In detail, the sensitivity analysis shows variability (median and 90% interval) in the predictive performance in a given environment (location, year combination) when we vary *i*) the hyperparameter values over their spesified ranges, *ii*) the genotype sets that we are predicting, and *iii*) the training set by removing any single training environment.

### Model

In the models *M*_*G+E*_, *M*_*G+E+GR*_ and 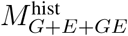 we assume that *i*) the yield *y*_*ij*_ of genotype *i* in environment *j* is affected by the genotype, the environmental conditions throughout the growing season and the interactions between the two. We assume that *ii)* the response to the environmental properties is non-linear and that *iii)* it may involve interactions between different environmental properties. For instance, temperature/rainfall either too low or too high reduces yield, and the response to rainfall is also affected by the soil type. We further assume that *iv)* the responses to the environmental conditions are highly polygenic. Assumptions *i-iv* are encoded using the kernel trick^15^, in which covariate data are represented as similarities, or kernels, between different data items. Kernel methods are a computationally effective way to model non-linearities and interactions and they have been applied to breeding data^16^. An additional complication in the data is the low number of observed trials compared to the complexity of the problem. To handle this, we constrain our model to only learn the most prominent combinations of environmental conditions affecting yield, by assuming a low-rank approximation for the model parameters accounting for the *G×E* effects. Finally, we follow the Bayesian statistical framework^17^, and regularise the model by placing priors on all parameters, which alleviates overfitting to the training data and improves prediction accuracy in the test data.

Mathematically, the model for yield is formulated as

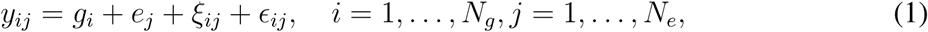

where *g*_*i*_ is the genetic main effect, *e*_*j*_ is the environmental effect, *ξ*_*ij*_ is the effect that arises from interaction between genotype *i* and environment *j*, *ϵ*_*ij*_ is noise distributed as 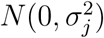, and *N*_*g*_ and *N*_*e*_ are the numbers of genotypes and environments. The genetic main effect *g*_*i*_ is modeled as a linear function of the genomic covariates. In detail, the model for the vector of genetic main effects g* = (*g*_1_,…, *g*_*N*_*g*__)^*T*^ is given in terms of a linear genomic kernel *K*_*g*_ by

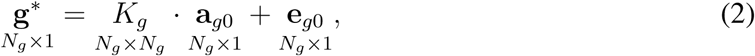

where a_*g0*_ are kernel regression weights and e_*g0*_ is the noise vector with elements distributed independently as 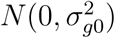. The dimension of each matrix is shown in equation (2) below the corresponding matrix symbol. The genomic kernel *K*_*g*_ is computed by first concatenating the genomic covariates *g*_*i*_ as the rows of a matrix G and then using the standard linear kernel, *K*_*g*_ = *GG*^*T*^.

The environmental main effect *e*_*j*_ in equation (1) is modeled as a random effect,

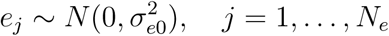

The *G×E* terms *ξ*_*ij*_ are modelled as non-linear functions of the genomic and environmental covariates, *g*_*i*_ and *e*_*j*_. Each environment and genotype is first represented by *R* latent variables. The interactions *ξ*_*ij*_ are modelled as the inner product of the latent variable vectors corresponding to genotype *i* and environment *j*, that is,

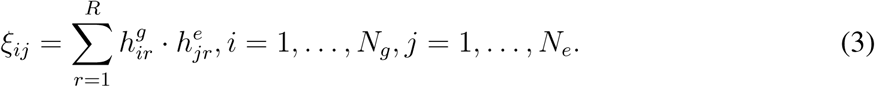

Here, 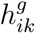 is the *k*th latent variable for the *i*th genotype, and 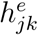 is the kth latent variable for the *j*th enviroment. Using matrix notation, equation (3) can be written as

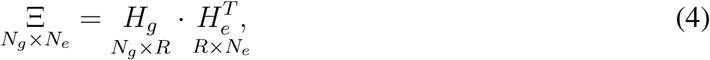

where Ξ = [*ξ*_*ij*_] is the matrix of interaction terms, and 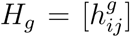 and the 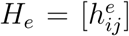 are matrices having as their rows the *R*-dimensional latent variable representations for each genotype and environment, respectively.

The latent variables *H*_*g*_ and *H*_*e*_ are obtained from genotype and environment kernels *K*_*g*_ and *K*_*e*_:

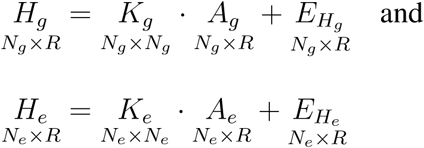

where *A*_*g*_ and *A*_*e*_ are kernel regression weights, and *E*_*H*_*g*__ and *E*_*H*_*e*__ are matrices containing error terms distributed independently as 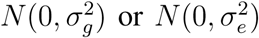, respectively. The environmental kernel *K*_*e*_ is obtained by combining multiple kernels 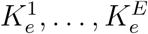, computed from enviromental data e_*j*_, *j* = 1,…,*N*_*e*_, each kernel representing a different aspect of the environment (weather, soil, etc). Details about processing the raw data into kernels and about combining multiple environmental kernels into a single kernel are presented in SI Data processing and kernels.

For inference we use variational approximation^18^, which is a computationally feasible way to approximate posterior distributions of parameters in complex models. The variational updates required here can be derived similarly to Gönen et al.^5^, except that we have extended their model and algorithm by including the genotype and environment main effects, i.e., the terms *g*_*i*_ and *e*_*j*_ in equation (1). Detailed distributions of the model parameters and the guidelines for specifying hyperparameter values are given in Sections SI Detailed model specification and SI Specifying hyperparameter values, respectively. Further details about the inference algorithm in SI Details of the variational inference algorithm.

### Comparison methods

The mixed model computations for the comparison methods `GBLUP` and `GE-BLUP` are performed using the **R** library `rrBLUP`^19^. For both methods, fixed effects were used to account for field block-specific effects, corresponding to the terms *e*_*j*_ in 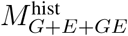, *M*_*G+E+GE*_ and *M*_*G+E*_. For `GBLUP`, the genomic kernel (see Section Model) was used as the covariance matrix Σ. For `GE-BLUP`, the environmental kinship model (GE-KE)^8^, is used and the full covariance matrix Σ is generated through the Kronecker product Σ = Σ_*G*_⊗ Σ_*E*_, where Σ_*G*_ and Σ_*E*_ are the genetic and environmental covariance matrices, respectively. The environmental covariance matrix Σ_*E*_ is generated from the available environmental data to describe soil properties and the growth conditions during the vegetative, heading time and grain filling developmental stages. All soil data and growth zone information are used as such whereas the daily average temperature and rainfall measurements are summarised as the mean and the standard deviation of the daily observations per crop stage. The growth periods are estimated using the sowing date and temperature sum accumulation-based estimates of heading and ripening times (440.2 °C and 905.9 °C, respectively), which were estimated from external breeding data. The vegetative stage is assumed to last 3 weeks starting from sowing, the time of heading is assumed to start 2 weeks before and last 1 week after the estimated heading time and grain filling was assumed to start after heading and to last 1 week longer than the estimated time of ripening. Wide estimates for the growth periods were used to account for varying growth speeds. The resulting set of environmental covariates is z-normalized and a linear kernel is used, which is further normalized according to equation (()) in SI Data preprocessing and kernels.

## Data availability

The data accompanied by the method code will be made available upon publication in the form of kernels to allow reproducing the results.

## Acknowledgements

We acknowledge Boreal Plant Breeding Ltd for the access to the plant breeding data and we gratefully thank Outi Manninen, Mika Isolahti and Esa Teperi for very helpful comments. This work was in part funded by Tekes, the Finnish Funding Agency for Innovation (Dnro 1718/31/2014 to J.G, H.M and S.K) and the Academy of Finland (Finnish Centre of Excellence in Computational Inference Research COIN, and grant numbers 294238, 292334 to SK and grant numbers 286607 and 294015 to P.M.).

## Author contributions

J.G. processed the data from Boreal Plant Breeding Ltd and performed the *in silico* experiments. J.G. and P.M. implemented the method. All authors were involved in the conception and design of the study, analyzed the results and assisted with drafting and critically revising the manuscript.

## Supplementary Information (SI)

### Data preprocessing and kernels

A summary of different kernels, including transformations specific to each data source, preprocessing and kernel transformations used, is given in Table S 1. The bandwidth parameter of all the Gaussian kernels is set to the conventional default value equal to the number of covariates used to compute the kernel. All kernels *K* are normalized to make them unit diagonal:

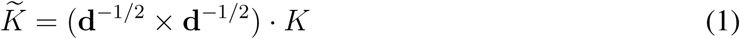

where d is a vector of the diagonal values of kernel *K*, × denotes the outer product, and the d^-1/2^ denotes a vector with all elements of d raised to the power of -1/2. The interaction kernel between the soil type and rainfall is computed from other kernels as

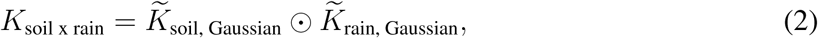

where ⊙ denotes the Hadamard (elementwise) product. Finally, all kernels are normalized with respect to their summed total variance by multiplication with a constant *c*

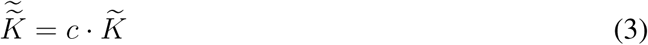

where 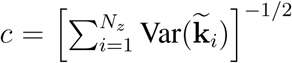 and 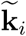 is the *i*th column of 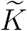. The motivation for this normalisation comes from the expectation that a priori each kernel explains the same amount of variance, and, when combining the kernels as described below, this prior expectation is realised by the normalisation.

### Combining environmental kernels

The final environmental kernel *K*_*e*_ is obtained as a weighted sum of the different normalized 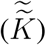 kernels in Table S 1. The weights are learnt from the training data by fitting **BEMKL**^20^, a multiple kernel learning regression method, using experiment-specific yield means as the target variable. For **BEMKL**, shape (*α*) and scale (*β*) parameter values for the prior Gamma distributions are set to 1 except for the *λ* parameter, for which the scale is fixed to 10, providing stronger regularization. Regression bias term *b* is set to 0. For further details of **BEMKL**, see Gönen et al.^20^. Before combining the kernels, the learnt weights are normalized such that their sum of squares is equal to 1, and the largest weight (in absolute value) is positive. The distributions of the normalised kernel weights from the sensitivity analysis are presented in Figure S 3. The composite kernel *K*_e_ is again normalized according to equation (3).

### Detailed model specification

The distributional assumptions of the model are

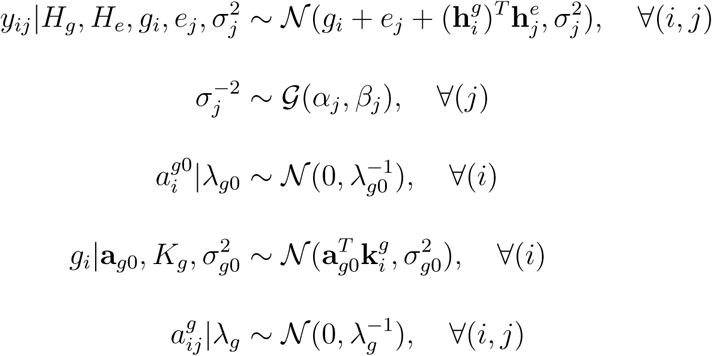

**Table S1.** Preprocessings and kernel functions applied to covariates.

| Variable<br>(unit) | transformation | preprocessing parameters | missing value imputation | kernel transformation(s) |
| --- | --- | --- | --- | --- |
| Soil content (% ,<br>$N_{\text{covs}} = 3$) | log transformation | z-normalization | (none) | linear and Gaussian |
| Soil organic content (% ,<br>$N_{\text{covs}} = 3$) | log transformation | z-normalization | (none) | Gaussian |
| daily rainfall (mm ,<br>$N_{\text{covs}} = 123$) | 7-day moving average (6 previous days) | z-normalization with 3<sup>rd</sup> order polynomial smoothing | 0-imputation | linear and Gaussian |
| daily average temperature (C° ,<br>$N_{\text{covs}} = 123$) | | z-normalization with 3<sup>rd</sup> order polynomial smoothing of daily mean/scale parameters | 0-imputation | linear and Gaussian |
| growth zone (1-4 ,<br>$N_{\text{covs}} = 1$) | | z-normalization | (none) | Gaussian |
| genotype markers (SNPs ,<br>$N_{\text{covs}} = 5696$) | | Minor allele frequency scaling for SNP A:<br>$\frac{A - 2 \cdot \text{MAF}_A}{\sqrt{2 \cdot \text{MAF}_A \cdot (1 - \text{MAF}_A)}}$ | mean imputation | linear kernel |

**Figure S3:**
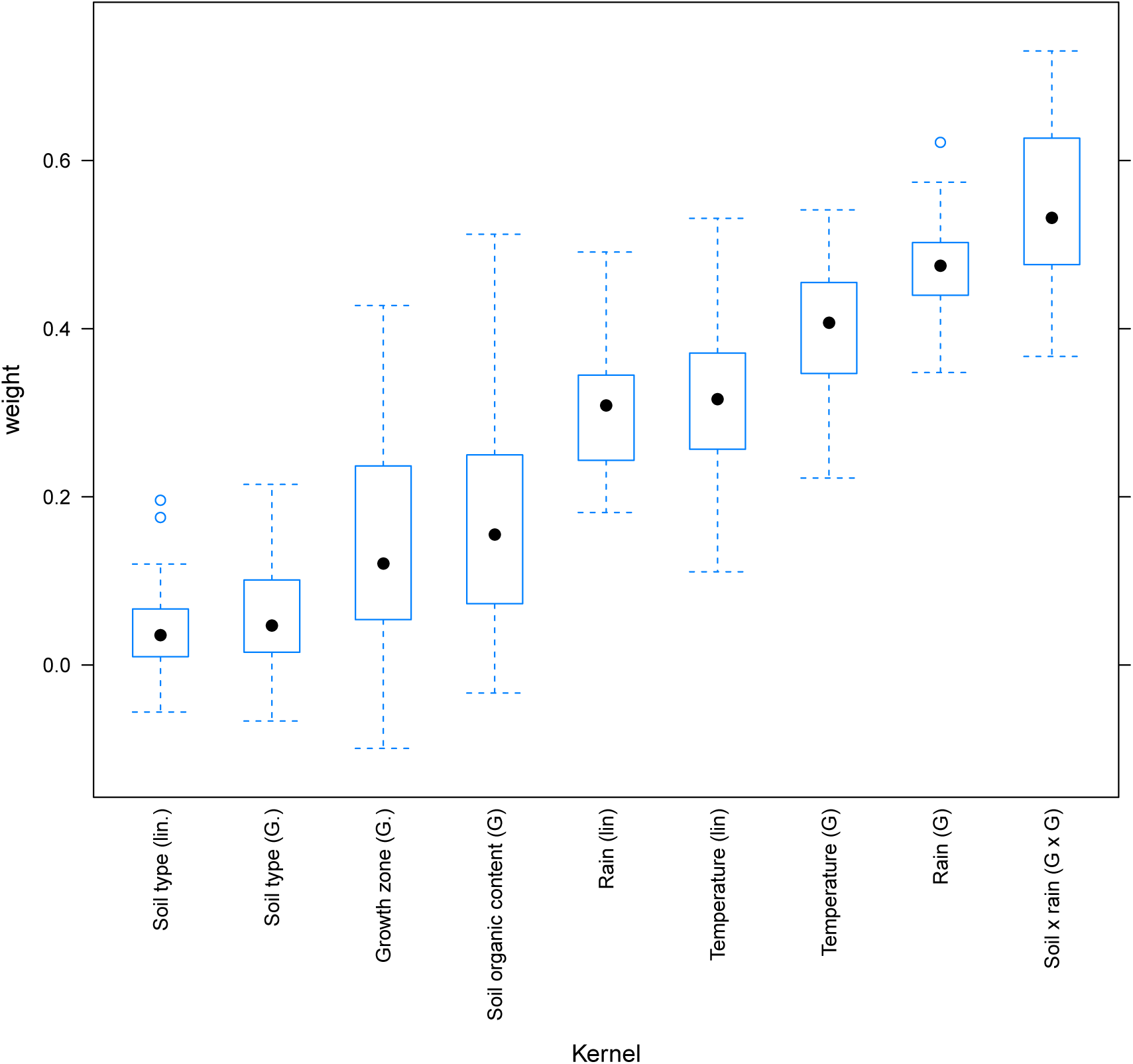
Estimated normalized kernel weights in the sensitivity analysis

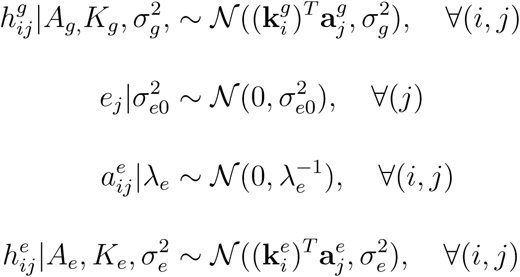

where 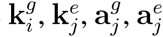, denote columns of matrices *K*_*g*_, *K*_*e*_, *A*_*g*_, *A*_*e*_, with subscripts *i* and *j* specifying the column index; 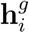 and 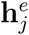 denote *i* th and *j*th rows of *H*_*g*_ and *H*_*e*_, represented as column vectors; 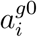 is the *i*th element of vector a_*g0*_; 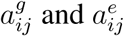 are the (*i, j*)th elements in matrices *A*_*g*_ and *A*_*e*_. 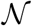 and 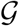 denote the Gaussian and Gamma distributions, respectively.

### Specifying hyperparameter values

Prior knowledge about the approximate weights of different sources of variance, e.g. the relative weight of genetic and environmental main effects, is used to specify hyperparameter values. We determine for each hyperparameter either a single fixed value or a grid of values to be selected from by cross-validation. Parameters (*α*_*j*_, *β*_*j*_) of the Gamma distribution for environment-specific residual noise variances 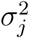 are set to (10, 1), corresponding to an expected value of approximately 0.1 for 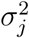. The variance of environment mean effects 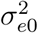 is fixed to 0.25. To set the parameters 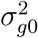 that determine the amount of signal and noise in the genetic main effects, we find values for them such that two conditions are satisfied. First, 95% of the variance of the genetic effects g* is assumed to be signal, that is,

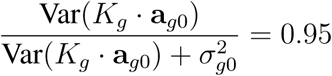

The second condition is that the variance of the genetic main effects, 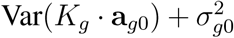, is either 0.2, 0.4, or 0.6. In practice we find these values for 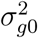 and λ_*g0*_ by simulating multiple realisations from the model with specific values for the parameters, and select values that on average satisfy the two conditions.

The parameters 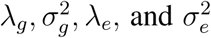, controlling the proportion of signal and noise in the latent components *H*_*g*_ and *H*_*e*_ that model the *G×E* interactions, are selected according to similar principles: by inspecting the proportion of signal of the total variance of the latent factors and the relative contribution of the interaction terms compared to the genetic main effects. In detail, we assume first that

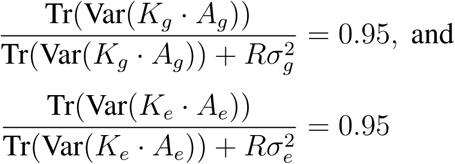

where Tr() denotes the trace of a matrix. Second, we assume that the total variance of the interactions is either the same or half of the total variance from the genetic main effects, i.e.

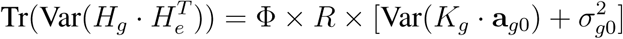

where Φ is either 0.5 or 1, to be selected with cross-validation.

### Details of the variational inference algorithm

For short-hand, the hyper-parameters in the model are denoted jointly by

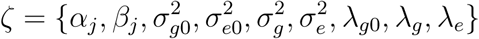

and the parameters by

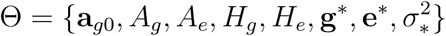

where 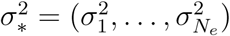. In the following the dependence on ζ is omitted for clarity. We assume the factorized variational approximation

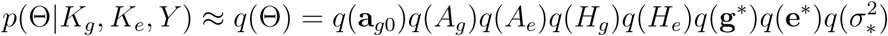

and define each factor in the ensemble just like its full conditional:

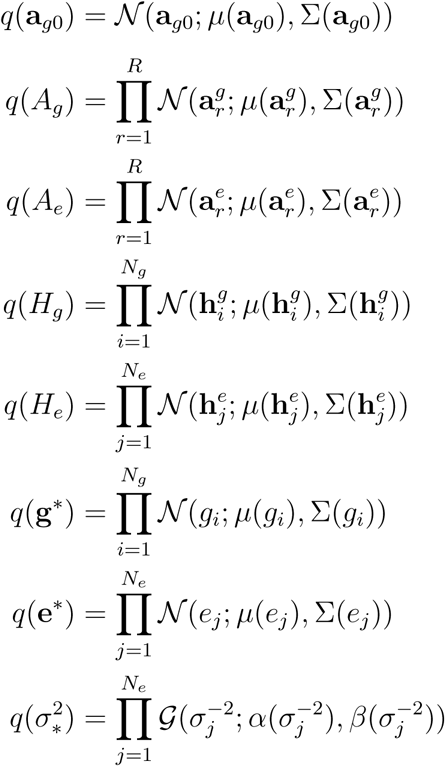

The parameters in the factor distributions can be derived as by Gönen et al^5^, and they are therefore omitted from here.

### Initialisation of the variational algorithm

The parameter g* was initialised to the main genetic effects learnt by `GBLUP`, and e* was initialised to the average yields in the different environments. Parameters *H*_*g*_ and *H*_*e*_ were initialised by applying the regularized Singular Value Decomposition (SVD) implemented in R library `softImpute` to the yield matrix *Y* after regressing out the initialised main effects g* and e*. Parameters a_*g0*_, *A*_*g*_ and *A*_*e*_ were initialised to 0. Environment-specific residual variance parameters 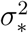 were initialised to environment-specific sample variances.

### Details of experimental setup

Different prediction tasks, distinguished by the availability of different data types, are presented in Figure S 4. Setups 1-4 correspond to those studied by Malosetti et al.^8^: in setup 1, phenotype measurements are available for the genotypes and environments to be predicted, and both genotypes and environmental covariates are fully observed. In setups 2 and 3, phenotype measurements are still available but only for the genotypes or the environments to be predicted, but not both, and covariates are fully observed. In setup 4, no phenotype data are available for environments/genotypes to be predicted, but both genetic and environmental covariates are still fully observed.

Two additional setups can be considered. In setups 5 and 6 environmental covariates from the environments of interest are only partially available: location and soil characteristics are known but the in-season weather measurements are not available for the year of interest. However, historical observations for the same locations are available and they are used to estimate the performance of each genotype. Setups 5 and 6 differ depending on whether phenotype measurements are available from some other environment for the genotypes (5) or not at all (6). The results in this paper are for setup 6 where no phenotype data are available for any of the lines of interest. We emphasize that a further difference to earlier work^8^ is that we strictly require the test environments to be simultaneously both from a location and from a year not included among the training environments and that the genotypes in the test/validation sets are from the progeny of the training set. A summary of the differences between our setup to those presented by earlier works is given in Table S2

**Figure S4:**
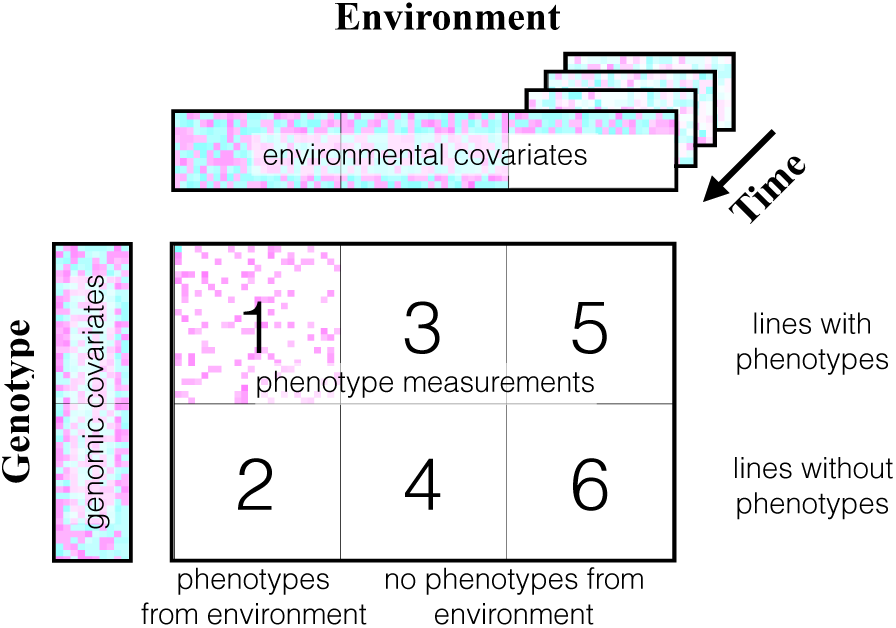
Comparison of prediction setups with respect to the availability of phenotype data and the genomic and environmental covariates as presented by Malosetti et al^8^. White colour indicates missing value. In setups 1, 3 and 5, ’’lines with phenotypes”, the lines to be predicted have phenotype observations (from some environments). In setups 1 and 2, “phenotypes from environment”, phenotypes have been measured from the prediction target environments (for some lines). In setups 1-4 presented by Malosetti et al^8^., environmental covariates are available for all environments, whereas in the new setups 5 and 6, environmental covariates from the trials of interest are missing and they are replaced by using several years of historical data.

### Gains from modelling *G×E* for current target population of environments

Our results indicate targeted breeding could improve yields by dividing a single target population of environments (TPE) into several parts, but the same methodology could be used even when developing only 1 variety for a larger population of target environments as in traditional breeding. Traditional breeding makes the implicit assumption that varieties’ observed yields *g* ∈1,…,*G* in trial experiments in environments (location × year) *e*ε1,…, *E*, are representative of the yield in the TPE, in other words

**Table S2:** Comparison of the proposed in *silico* setup to the existing setups.

| Publication | New environment | New genotypes |
| --- | --- | --- |
| Burgueño et al. 2012 (CV1/CV2)<sup>10</sup> | CV1/CV2: test locations and years are present in the location-year combinations in the training data | new lines in CV1: not restricted to the offspring generation. In CV2 the test lines have phenotype observations |
| Heslot et al. 2013<sup>7</sup> | Random split, balanced wrt years and locs → year and locations not new | only 544/2195 genotypes have no phenotype observations, test set not restricted to the offspring generation |
| Albrecht et al. 2014<sup>11</sup> | the year-location combination is new but the test locations and years are present in other location-year combinations in the training data | genotypes are new and from the offspring |
| Malosetti et al. 2016<sup>8</sup> | time-structured DTD: 2/6 test locations new according to strict criteria; physically structured DTD: none of the environments are strictly new (as the year is not new) | all genotypes within the same family, not from the next generation. |
| Saint Pierre et al. 2016<sup>12</sup> (leave-one-side-out) | location new but year part of the training set | test lines have phenotype observations |

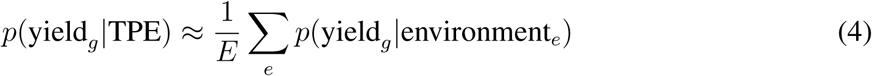

However, with geographic field use information and weather data widely available, this strong assumption can be replaced with an estimate for the yield in the TPE given the actual fields and their microclimates:

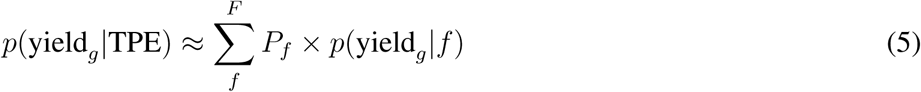

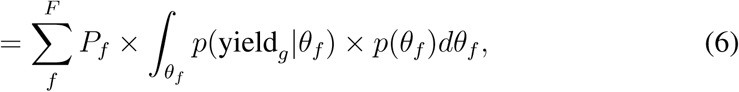

where *f* ∈ 1,…,*F*, are fields in the TPE used for cultivation of the new variety, *θ*_*f*_ are parameters (e.g. weather conditions) related to a certain field *f*, *p* (*θ*_*f*_) is the uncertainty related to these conditions, estimated from historical records, *p* (yield_*g*_|*θ*_*f*_) is the predictive distribution for the yield under conditions *θ*_*f*_, obtained from the model, and *P*_*f*_ is the proportion of the total volume cultivated in field *f*.

**Table S3:**
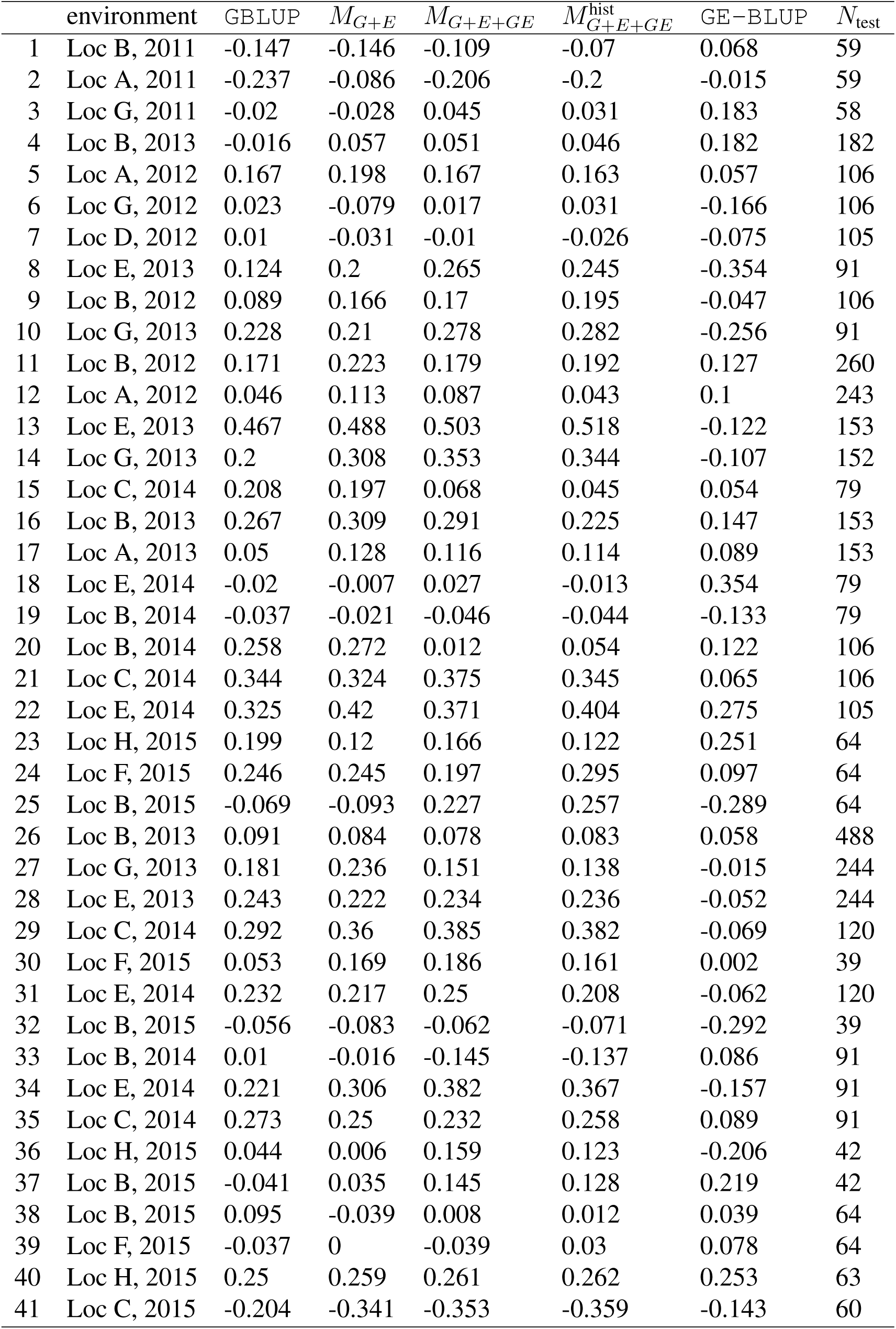
Results for individual test folds.

